# *flt1* inactivation promotes zebrafish cardiac regeneration by enhancing endothelial activity and limiting the fibrotic response

**DOI:** 10.1101/2024.09.11.612516

**Authors:** Zhen-Yu Wang, Armaan Mehra, Qian-Chen Wang, Savita Gupta, Agatha Ribeiro da Silva, Thomas Juan, Stefan Günther, Mario Looso, Jan Detleffsen, Didier Y.R. Stainier, Rubén Marín-Juez

## Abstract

**Summary statement:** *flt1* inactivation promotes zebrafish cardiac regeneration by enhancing coronary revascularization and endocardial expansion, while limiting myofibroblast differentiation.

VEGFA administration has been explored as a pro-angiogenic therapy for cardiovascular diseases including heart failure for several years, but with little success. Here we investigate a different approach to augment VEGFA bioavailability: by deleting the VEGFA decoy receptor VEGFR1/FLT1, one can achieve more physiological VEGFA concentrations. We find that following cryoinjury, zebrafish *flt1* mutant hearts display enhanced coronary revascularization and endocardial expansion, increased cardiomyocyte dedifferentiation and proliferation, and decreased scarring. Suppressing Vegfa signaling in *flt1* mutants abrogates these beneficial effects of *flt1* deletion. Transcriptomic analyses of cryoinjured *flt1* mutant hearts reveal enhanced endothelial MAPK/ERK signaling and downregulation of the transcription factor gene *egr3*. Using newly generated genetic tools, we observe *egr3* upregulation in the regenerating endocardium, and find that Egr3 promotes myofibroblast differentiation. These data indicate that with enhanced Vegfa bioavailability, the endocardium limits myofibroblast differentiation via *egr3* downregulation, thereby providing a more permissive microenvironment for cardiomyocyte replenishment after injury.

## Introduction

Due to the limited regenerative capacity of the adult human heart, myocardial infarction (MI) causes permanent loss of myocardium and fibrotic remodeling, ultimately leading to heart failure (Laflamme and Murry, 2011; McMurray and Pfeffer, 2005; Murry et al., 2006; Senyo et al., 2013; Tanai and Frantz, 2015). The formation of new blood vessels following MI is crucial for the survival and function of cardiac tissue. Achieving adequate revascularization has been a primary goal in the field of regenerative therapy for heart disease. However, attempts to promote revascularization and cardiac regeneration by administering pro-angiogenic factors such as vascular endothelial growth factor A (VEGFA) have thus far had limited success (Lupu et al., 2020).

Contrary to humans, adult zebrafish possess a remarkable ability to regenerate their heart wherein lost cardiomyocytes are replenished and fibrosis is resolved following injury (Bevan et al., 2020; Chablais et al., 2011; Gonzalez-Rosa et al., 2011; Poss et al., 2002). After injury, the zebrafish heart has the innate ability to efficiently revascularize damaged tissues (El-Sammak et al., 2022; Marin-Juez et al., 2019; Marin-Juez et al., 2016). Vegfaa regulates coronary revascularization in zebrafish hearts by promoting intra-ventricular vessel sprouting, which, along with superficial sprouting, creates a supportive framework for newly regenerated cardiomyocytes to occupy the injured tissue (Marin-Juez et al., 2019). VEGFA primarily interacts with two transmembrane tyrosine kinases predominantly expressed in endothelial cells: VEGF receptor 1 (VEGFR1/FLT1) and VEGFR2 (de Vries et al., 1992; Millauer et al., 1993). While VEGFR2 is recognized as the primary VEGFA signaling receptor due to its robust kinase activity following binding, FLT1 binds VEGFA with even greater affinity (Roberts et al., 2004), but induces very weak kinase activity (Ito et al., 1998; Waltenberger et al., 1994). Besides the transmembrane form, *FLT1* also encodes a soluble form (sFLT1) that lacks its transmembrane and kinase domains, rendering it incapable of inducing kinase activity even when bound to VEGFA (Shibuya, 2013). Therefore, FLT1 is usually regarded as a decoy receptor that attenuates VEGFR2 signaling by sequestering VEGFA. Previous studies have shown that *Vegfr1^-/-^* mice display embryonic lethality (Fong et al., 1995), while mice lacking *Vegfr1* function in endothelial cells exhibit hypervascularization as well as angiogenesis-induced cardiomyocyte growth (Kivela et al., 2019).

Previous studies in zebrafish have shown that constitutive overexpression of Vegfaa can stimulate cardiomyocyte cell cycle re-entry while blocking cardiac regeneration, further highlighting the importance of dosage and timing (Karra et al., 2018). More recently, a dose dependent mitogenic effect of VEGFA has been reported (Pontes-Quero et al., 2019). We hypothesize that more physiological levels of VEGFA promote cardiac regeneration. To test this hypothesis and identify pro-regenerative mechanisms, we investigate cardiac regeneration in *flt1* mutant zebrafish.

*flt1* deletion in zebrafish enhances the response of the cardiac endothelium to injury, and also enhances cardiomyocyte regeneration and reduces scarring. Furthermore, we find that *flt1* deletion downregulates the transcription factor gene *egr3*. Manipulation of *egr3* levels during cardiac regeneration reveals a role for Egr3 in promoting myofibroblast differentiation. Overall, our data indicate that physiological stimulation of Vegfa signaling accelerates cardiac regeneration by enhancing endothelial replenishment and limiting myofibroblast differentiation and reducing scarring. These conditions might create a more permissive milieu for cardiomyocyte regeneration.

## Results

### *flt1* deletion promotes coronary vascular development and endothelial regeneration after cardiac cryoinjury

Previous studies have shown that in zebrafish larvae, *flt1* is expressed predominantly in endothelial cells and that it plays a pivotal role in modulating angiogenesis and vessel branching morphogenesis (Krueger et al., 2011; Zygmunt et al., 2011). While disrupting the signaling function of the membrane form of *flt1* in zebrafish has no obvious effect on angiogenesis, mutations affecting the function of its soluble form enhance endothelial growth in a Vegfaa dependent manner (Matsuoka et al., 2016; Wild et al., 2017). Therefore, we hypothesized that Flt1 might also be involved in limiting coronary vessel growth during development and regeneration. To test this hypothesis, we utilized the *Tg(−0.8flt1:RFP)* line to visualize coronary endothelial cells (cECs) and crossed it with *flt1* mutants and a *Tg(hsp70l:sflt1)* overexpression line. Coronary formation begins 1-2 months after fertilization (Harrison et al., 2015). We analysed coronary network expansion at 42 days post fertilization (dpf) (body length ∼ 20 mm), a timepoint when coronary vessels begin to form a basic network in wild-type hearts. To ensure a sustained *sflt1* overexpression during the initiation of coronary network formation, we implemented a daily heat shock treatment from 28 to 42 dpf. We found that coronary network expansion in *flt1* mutants was enhanced, observing increased branching and vascular coverage over the ventricles (Fig. S1A-B). In contrast, the formation of the coronary network was notably suppressed in juvenile hearts overexpressing *sflt1*, observing only a few sprouts around the atrioventricular canal (Fig. S1C-D). While unaffected in *flt1* mutants, ventricular volume was significantly reduced after developmental overexpression of *sflt1* (Fig. S1E). When analyzing ventricles at adult stages, we found no differences in vessel coverage between *flt1^-/-^* and *flt1^+/+^* siblings (Fig. S1F-G).

Next, we set out to test whether Flt1 regulates coronary regeneration. To this end, we cryoinjured the ventricles of *flt1* mutants and *Tg(hsp70l:sflt1)* fish in the *Tg(−0.8flt1:RFP)* background. For the *Tg(hsp70l:sflt1)* experiments, we implemented daily heat shock treatments before and after the cryoinjury until the observation timepoints (Fig. S2A), as described in our previous study (El-Sammak et al., 2022). We analysed coronary coverage of the injured tissue in *flt1* mutants at 96 hours post cryoinjury (hpci), when revascularization of the injury is obvious and coronary endothelial cell proliferation peaks in wild types (Marin-Juez et al., 2016; Ross Stewart et al., 2022). We found that *flt1* mutants exhibited significantly enhanced revascularization of the injured tissue compared with wild types as measured by coronary vessel coverage (Fig. 1A-A”). In contrast, zebrafish overexpressing *sflt1* exhibited significantly impaired revascularization at 7 days post cryoinjury (dpci) (Fig. 1B-B”), a timepoint when regenerating coronaries fully cover the injured tissue in wild types (El-Sammak et al., 2022). Injured areas in *sflt1* overexpressing zebrafish were still un-revascularized at 30 and 90 dpci (Fig. 1C-D’). To look more closely at the cECs, we quantified their proliferation in the border zone and injured area (BZI) at 96 hpci and found that it was significantly increased in *flt1* mutants and decreased in *sflt1* overexpressing zebrafish when compared with wild-type and control siblings, respectively (Fig. 1E-H).

**Fig. 1.**
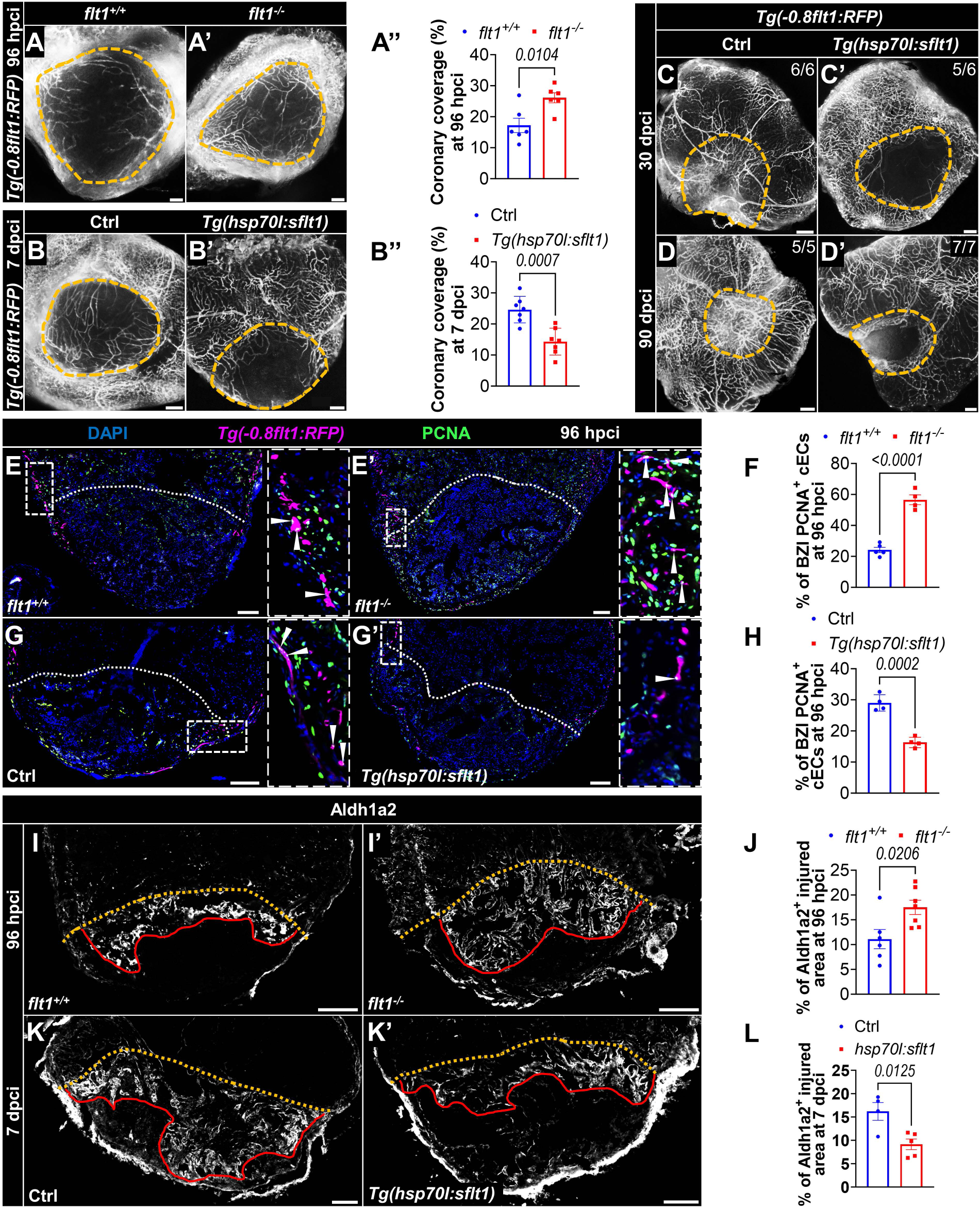
Flt1 regulates endothelial regeneration. **(A-B”)** Wholemount images of cryoinjured ventricles from *Tg(−0.8flt1:RFP);flt1^+/+^* (n=6, A) and *Tg(−0.8flt1:RFP);flt1^-/-^*(n=6, A’) zebrafish, and from *Tg(−0.8flt1:RFP)* (Ctrl, n=7, B) and *Tg(− 0.8flt1:RFP);Tg(hsp70l:sflt1)* (n=7, B’) zebrafish showing revascularization of injured area at 96 hpci and 7 dpci respectively, and the corresponding percentage of coronary coverage of the injured area (A”,B”). **(C-D’)** Wholemount images of cryoinjured ventricles from *Tg(− 0.8flt1:RFP)* (Ctrl, C,D) and *Tg(−0.8flt1:RFP);Tg(hsp70l:sflt1)* (C’,D’) zebrafish showing revascularization at 30 and 90 dpci. **(E,E’,G,G’)** Immunostaining for RFP (cECs, magenta) and PCNA (proliferation marker, green) with DAPI (blue) counterstaining on sections of cryoinjured ventricles from *Tg(−0.8flt1:RFP);flt1^+/+^* (n=5, E) and *Tg(−0.8flt1:RFP);flt1^-/-^* (n=4, E’) zebrafish, and from *Tg(−0.8flt1:RFP)* (Ctrl, n=4, G) and *Tg(−0.8flt1:RFP);Tg(hsp70l:sflt1)* (n=4, G’) zebrafish at 96 hpci. Arrowheads point to PCNA^+^ cECs. **(F,H)** Percentage of PCNA^+^ cECs in the BZI of the indicated genotypes at 96 hpci. **(I,I’,K,K’)** Immunostaining for Aldh1a2 (activated endocardium, white) on sections of cryoinjured ventricles from *flt1^+/+^*(n=6, I) and *flt1^-/-^* (n=7, I’) zebrafish at 96 hpci, and from non-transgenic Ctrl (n=4, K) and *Tg(hsp70l:sflt1)* (n=5, K’) zebrafish at 7 dpci. Red lines demarcate the extent of activated endocardium in the injured area. **(J,L)** Percentage of Aldh1a2^+^ injured area from the indicated genotypes. Orange and white dotted lines demarcate the injured areas. White dotted boxes correspond to the magnified regions. Scale bars: 100 µm. Data show mean ± SEM (two-tailed, unpaired Student’s *t*-test with *p* values shown in the graphs).

FLT1 serves as a decoy receptor for VEGFA thereby reducing VEGFA bioavailability for VEGFR2 and attenuating VEGFA/VEGFR2-induced angiogenesis (Ho et al., 2012; Kivela et al., 2014; Robciuc et al., 2016; Ruiz de Almodovar et al., 2009). Indeed, the pro-angiogenic phenotype in *flt1* mutants has been shown to be due to increased Vegfa bioavailability (Matsuoka et al., 2016; Wild et al., 2017). To further confirm that the observed phenotypes are Vegf dependent, we overexpressed a dominant-negative form of *vegfaa* (*dnvegfaa*) (Muller et al., 1997a; Muller et al., 1997b; Rossi et al., 2016) in *flt1* mutants using the *Tg(hsp70l:dnvegfaa)* line (Marin-Juez et al., 2016) and quantified coronary coverage of the injured area and cEC proliferation at 96 hpci. Overexpression of *dnvegfaa* in *flt1* mutants blocked most of the increase in tissue revascularization and cEC proliferation observed after *flt1* deletion (Fig. S3A-D).

The observed phenotypes in cECs prompted us to investigate whether modulation of *flt1* affects the behavior of the other cardiac endothelial cell population, the endocardium, given that revascularization is partially regulated by endocardial cues (Marin-Juez et al., 2019). We utilized Aldh1a2 as a marker to identify activated endocardial cells (EdCs)(Kikuchi et al., 2011; Munch et al., 2017) and found that *flt1* mutants display increased expansion of Aldh1a2^+^ cells within the injured area compared to wild types at 96 hpci (Fig. 1I-J). This finding was further corroborated by co-staining for Aldh1a2 and Fli1 and quantifying the coverage of Aldh1a2^+^/Fli1^+^ cells within the injured area (Fig. S2B-C). Given that EdC proliferation is high at 96 hpci (Munch et al., 2017), we also assessed it at this timepoint and found a significant increase within the injured area (Fig. S2D-E). While *sflt1* overexpression strongly reduced cEC proliferation (Fig. 1G-H), it did not appear to affect endocardial expansion at 96 hpci (Fig. S2F-G). Interestingly, *sflt1* overexpression reduced endocardial expansion at 7 dpci (Fig. 1K-L). Altogether, these data indicate that Vegfa availability determines coronary regeneration after cardiac cryoinjury in zebrafish. Moreover, increased Vegfa signaling can enhance endocardial expansion.

### *flt1* modulation alters cardiomyocyte regeneration and scarring after cardiac cryoinjury

During cardiac regeneration, cardiomyocytes undergo dedifferentiation and proliferation (Beisaw et al., 2020; Kikuchi et al., 2010; Morikawa et al., 2015; Tsedeke et al., 2021), processes regulated, at least in part, by cECs (Marin-Juez et al., 2019; Marin-Juez et al., 2016) and EdCs (Galvez-Santisteban et al., 2019; Kikuchi et al., 2011; Munch et al., 2017; Zhao et al., 2019). In view of the endothelial phenotypes observed upon manipulation of *flt1* expression, we sought to investigate the impact on cardiomyocyte regeneration. Using the embryonic myosin heavy chain antibody N2.261 as a readout for cardiomyocyte dedifferentiation (Sallin et al., 2015), and the DNA replication marker PCNA as an indicator for cardiomyocyte proliferation, we assessed the percentage of border zone dedifferentiating and proliferating cardiomyocytes in *flt1*^-/-^ and *Tg(hsp70l:sflt1)* zebrafish after cardiac cryoinjury. We observed a marked increase in both cardiomyocyte dedifferentiation and proliferation in *flt1* mutants at 96 hpci and 7 dpci (Fig. S3A-D, Fig. 2A-D). Notably, cardiomyocyte proliferation in *flt1* mutants at 96 hpci was comparable to that observed in *flt1* mutants at 7 dpci (Fig. S4C-D and Fig. 2D), when cardiomyocyte proliferation in wild types is at its highest, suggesting a shift in cardiomyocyte proliferation dynamics. In contrast, cardiomyocyte dedifferentiation and proliferation were significantly reduced in *sflt1* overexpressing zebrafish when compared with controls (Fig. 2E-H). Together, these results suggest that *flt1* negatively regulates cardiomyocyte dedifferentiation and proliferation during cardiac regeneration. Additionally, we assessed cardiomyocyte behavior in *flt1* mutants overexpressing *dnvegfaa* and found that the increase in cardiomyocyte proliferation was also blocked (Fig. S3E-F).

**Fig. 2.**
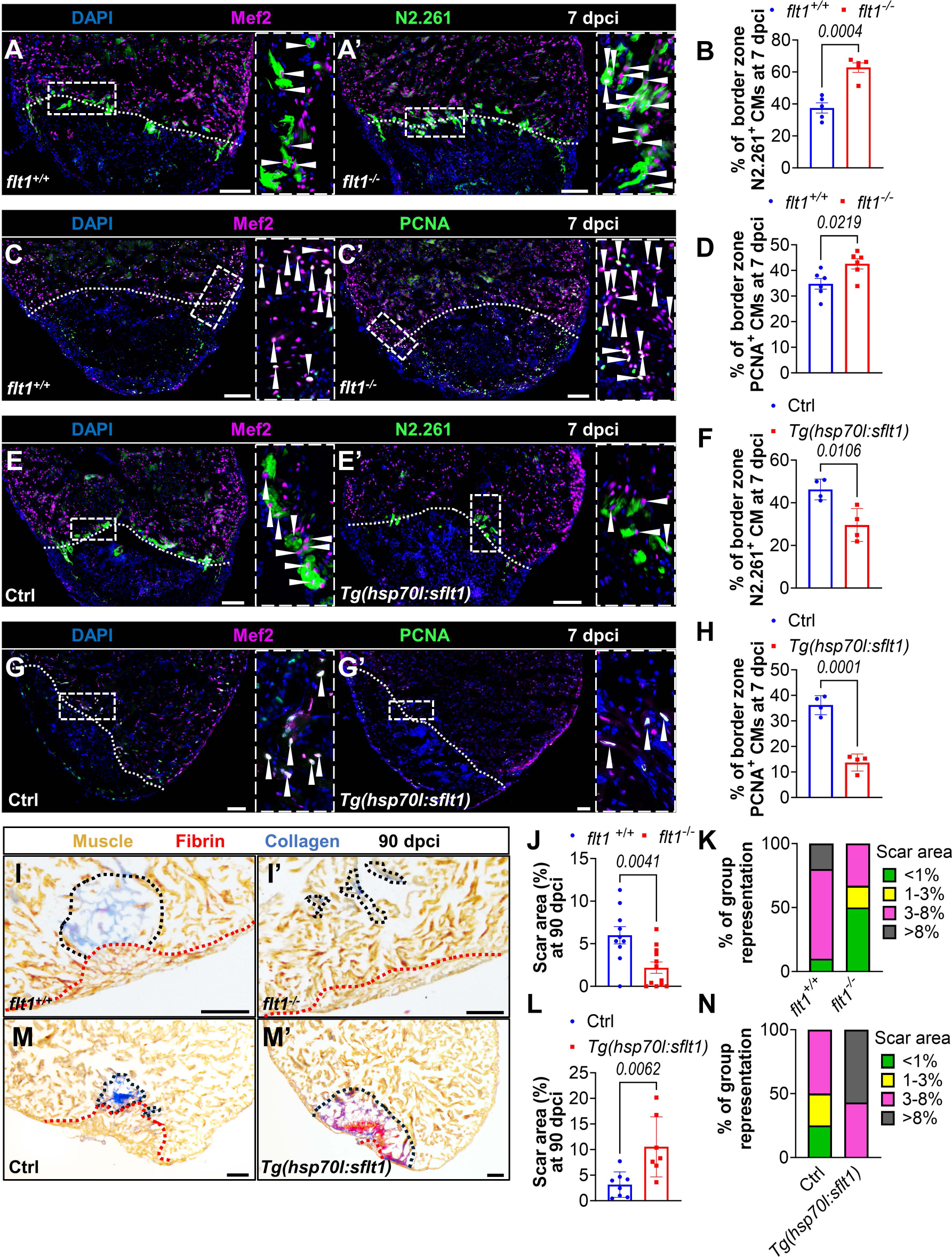
*flt1* modulation alters cardiomyocyte regeneration and scarring after cardiac cryoinjury. **(A,A’,E,E’)** Immunostaining for Mef2 (CM nuclei, magenta) and N2.261 (embryonic myosin heavy chain, green) with DAPI (blue) counterstaining on sections of cryoinjured ventricles from *flt1^+/+^* (n=5, A) and *flt1^-/-^*(n=5, A’) zebrafish, and from non-transgenic Ctrl (n=4, E) and *Tg(hsp70l:sflt1)* (n=4, E’) zebrafish at 7 dpci. Arrowheads point to N2.261^+^ CMs. **(B,F)** Percentage of N2.261^+^ CMs in the border zone of the indicated genotypes at 7 dpci. **(C,C’,G,G’)** Immunostaining for Mef2 (magenta) and PCNA (green) with DAPI (blue) counterstaining on sections of cryoinjured ventricles from *flt1^+/+^*(n=6, C) and *flt1^-/-^* (n=6, C’) zebrafish, and from non-transgenic Ctrl (n=4, G) and *Tg(hsp70l:sflt1)* (n=4, G’) zebrafish at 7 dpci. Arrowheads point to PCNA^+^ CMs. **(D,H)** Percentage of PCNA^+^ CMs in the border zone of the indicated genotypes at 7 dpci. **(I-I’,M-M’)** AFOG staining on sections of cryoinjured ventricles from *flt1^+/+^* (n=10, I) and *flt1^-/-^* (n=12, I’) zebrafish, and from non-transgenic Ctrl (n=8, M) and *Tg(hsp70l:sflt1)* (n=7, M’) zebrafish at 90 dpci. Orange, muscle; red, fibrin; blue, collagen. Black and red dotted lines delineate scar and regenerated muscle wall areas, respectively. **(J,L)** Percentage of scar area relative to ventricular area in of the indicated genotypes at 90 dpci. **(K,N)** Graphs showing the representation of groups of different scar area sizes of the indicated genotypes at 90 dpci. White dotted lines demarcate the injured areas. White dotted boxes correspond to the magnified regions. Scale bars: 100 µm. Data show mean ± SEM (two-tailed, unpaired Student’s *t*-test with *p* values shown in the graphs).

Given the impact of *flt1* modulation on cardiomyocyte regeneration, we further explored whether these alterations affect scarring during cardiac regeneration. In line with the other phenotypes, we observed enhanced cardiac regeneration at 90 dpci in *flt1* mutants, characterized by significantly smaller scar areas when compared with wild types (Fig. 2I-K). Conversely, ventricles overexpressing *sflt1* failed to regenerate, as evidenced by the presence of fibrin-rich large scars at both 30 and 90 dpci (Fig. S4E-G, Fig. 2M-N), consistent with observations from wholemount samples (Fig. 1C’,D’). In contrast, wounds in control cryoinjured ventricles displayed continuous myocardial wall enclosing the injury with collagen-rich scars at 30 dpci, and limited scar area at 90 dpci (Fig. S4E, Fig. 2M). These findings collectively indicate that *flt1* limits myocardial renewal and increases scarring during cardiac regeneration.

### *flt1* deletion causes alterations in endothelial MAPK/ERK signaling and downregulation of *egr3* during cardiac regeneration

To gain further insight into how *flt1* regulates cardiac regeneration, we conducted transcriptional analysis of the BZI from *flt1^+/+^* and *flt1^-/-^* sibling zebrafish at 96 hpci (Fig. S5A). We identified 85 differentially expressed genes (DEG) (*padj* < 0.05), with 72 of them showing downregulation in *flt1* mutants (Fig. S5B,C). Gene Ontology and KEGG pathway analyses reveal the significant downregulation of genes associated with negative regulation of MAPK/ERK signaling (Fig. S5D). Further investigation of the DEG within these categories revealed a decrease in the expression of genes encoding MAPK/ERK antagonists including *spry4*, *dusp1*, *dusp4*, and *dusp6* in cryoinjured *flt1* mutant ventricles, with the decreased expression of multiple anti-proliferative factor genes such as *btg2*, *gadd45ga*, and *igfbp1a*. Additionally, we observed an upregulation of the angiogenic factor gene *aplnra* among the few upregulated genes in cryoinjured *flt1* mutant ventricles. Changes in expression levels of these genes were confirmed by RT-qPCR analysis (Fig. 3A,B).

**Fig. 3.**
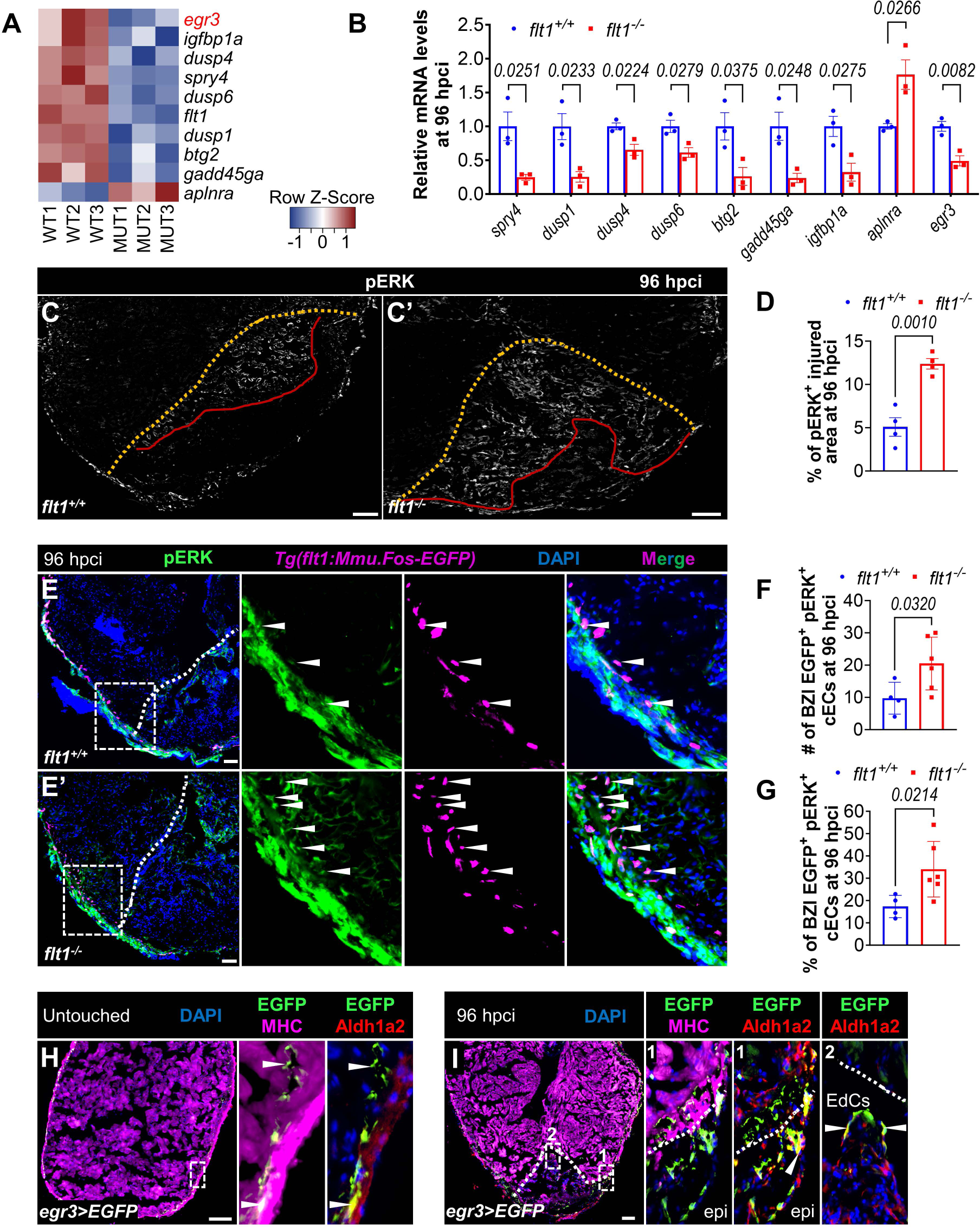
*flt1* deletion causes alterations in endothelial MAPK/ERK signaling and downregulation of *egr3* during cardiac regeneration. **(A,B)** Heat-map (A) and RT-qPCR validation (B) showing differentially regulated genes of interest in the BZI of cryoinjured ventricles from *flt1^+/+^*and *flt1^-/-^* zebrafish at 96 hpci. **(C,C’)** Immunostaining of cryoinjured ventricles from *flt1^+/+^*(n=4, C) and *flt1^-/-^* (n=4, C’) zebrafish at 96 hpci for pERK (phosphorylated ERK, white). Red lines demarcate the range of pERK^+^ injured area. **(D)** Quantification of the percentage of pERK^+^ injured area of the indicated genotypes at 96 hpci. **(E,E’)** Immunostaining for pERK (green) and EGFP (cECs, magenta) with DAPI (blue) counterstaining on sections of cryoinjured ventricles from *Tg(flt1:Mmu.Fos-EGFP);flt1^+/+^* (n=4, E) and *Tg(flt1:Mmu.Fos-EGFP);flt1^-/-^*(n=6, E’) zebrafish at 96 hpci. Arrowheads point to pERK^+^ cECs in the BZI. **(F,G)** Quantification showing the number (F) and percentage (G) of pERK^+^ cECs in the BZI of the indicated genotypes at 96 hpci. **(H,I)** Immunostaining for EGFP (green), MHC (myosin heavy chain, magenta), and Aldh1a2 (red) with DAPI (blue) counterstaining on representative sections from untouched (H) and cryoinjured (I) *egr3>EGFP* ventricles at 96 hpci. Arrowheads in (H) point to EGFP*^+^* cells. Orange and white dotted lines demarcate the injured areas. White dotted boxes correspond to the magnified regions. epi, epicardium. Scale bars: 100 µm. Data show mean ± SEM (two-tailed, unpaired Student’s *t*-test with *p* values shown in the graphs).

Previous studies have revealed a role for MAPK/ERK signaling in zebrafish cardiac regeneration, and its activation in the injured endocardium (Cardeira-da-Silva et al., 2024; Liu and Zhong, 2017; Tahara et al., 2021). We speculated that the downregulation of MAPK/ERK antagonists might facilitate MAPK/ERK activation in endocardial cells following cardiac cryoinjury. After injury, MAPK/ERK signaling is enhanced in the activated endocardium as assessed by phospho-ERK (pERK) immunostaining (Fig. S5E) (Cardeira-da-Silva et al., 2024). Notably, we observed a significant expansion of pERK^+^ endocardium within the injured area in *flt1* mutants compared with wild types (Fig. 3C-D). By using the *Tg(flt1:Mmu.Fos-EGFP)* line (El-Sammak et al., 2022) to label specifically the coronary endothelium, we also observed a significant increase in both the number and percentage of pERK^+^ coronary vessels at the border zone in cryoinjured *flt1* mutant ventricles (Fig. 3E-G’). Additionally, *flt1* mutants exhibited a higher abundance of coronary vessels at the border zone (Fig. 3E’), consistent with a previous report (Marin-Juez et al., 2019). Taken together, these findings indicate that *flt1* deletion enhances endothelial MAPK/ERK signaling following cardiac cryoinjury.

Interestingly, one of the most downregulated genes in cryoinjured *flt1* mutant ventricles is *early growth response 3* (*egr3*) (Fig. 3A,B, Fig. S5C), a MAPK/ERK signaling effector gene (Herndon et al., 2014; Li et al., 2007). To determine the expression pattern of *egr3* after cardiac cryoinjury, we utilized the recently generated knock-in *egr3* reporter line *Pt(egr3:Gal4-VP16);Tg(5xUAS:EGFP)* (da Silva et al., 2024), abbreviated as *egr3>EGFP*. Immunostaining for GFP, MHC, and Aldh1a2 revealed that in untouched ventricles, *egr3>EGFP* is only marginally expressed between the cortical and trabecular myocardial layers, as well as in the epicardium (Fig. 3H). After cryoinjury, *egr3>EGFP* expression was induced in the injured area, characterized by broader expression within and alongside epicardium-derived cells (EPDCs) covering the injured tissue and with expression in the Aldh1a2^+^ EdCs expanding into the wound (Fig. 3I).

### *flt1* deletion limits myofibroblast differentiation and promotes cardiomyocyte repopulation by downregulating *egr3*

We noted that the *egr3>EGFP* expression pattern in both untouched and cryoinjured ventricles closely resembles the distribution of fibroblasts (Sanchez-Iranzo et al., 2018). To better define the different cell types expressing *egr3* before and after cardiac cryoinjury, we analysed a previously published single-cell RNA sequencing dataset (Koth et al., 2020). This analysis revealed that following cryoinjury, *egr3* is upregulated in certain endothelial cell clusters, as well as in fibroblasts and myofibroblasts (Fig. S6A-D). We then conducted immunostaining on cryoinjured *egr3>EGFP* ventricles at 7 dpci for GFP, Aldh1a2, and α-SMA, and observed that about 36% of Aldh1a2^+^ EdCs expressed *egr3>EGFP*, with these EdCs being positive for α-SMA or located in close proximity to α-SMA^+^ cells (Fig. 4A,B). To further characterize these *egr3>EGFP*-expressing EdCs, we utilized Vimentin (Vim) as a marker for fibroblasts/myofibroblasts and found that they were indeed positive for Vim (Fig.4C). These data are indicative of their transition towards a fibroblast/myofibroblast identity, consistent with the reanalysis results of the single-cell RNA sequencing dataset. These results lead us to hypothesize that *egr3* promotes endothelial-to-mesenchymal transition (EndoMT) during zebrafish heart regeneration.

**Fig. 4.**
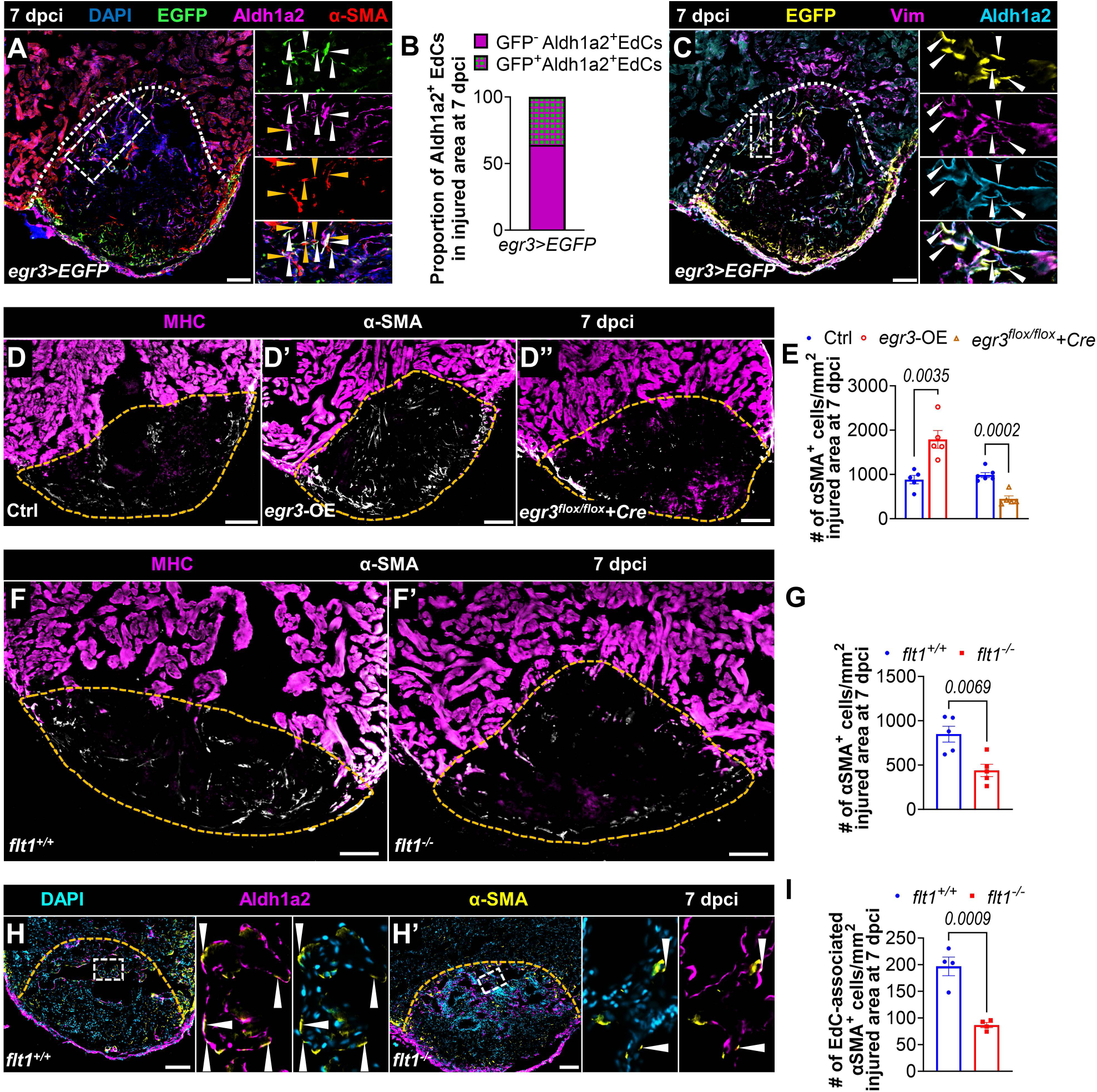
*flt1* deletion limits myofibroblast differentiation. **(A)** Immunostaining for EGFP (green), Aldh1a2 (magenta), and α-SMA (myofibroblast, red) with DAPI (blue) counterstaining on representative sections of cryoinjured *egr3>EGFP* ventricles at 7 dpci. White and orange arrowheads point to the EGFP^+^Aldh1a2^+^ EdCs and α-SMA^+^ cells, respectively. **(B)** Proportion of EGFP^+^ EdCs within the wound of cryoinjured *egr3>EGFP* ventricles (n=3). **(C)** Immunostaining for GFP (yellow), Aldh1a2 (cyan), and Vim (fibroblast/myofibroblast, magenta) with DAPI (blue) counterstaining on representative sections of cryoinjured *egr3>EGFP* ventricles at 7 dpci. Arrowheads point to the EGFP^+^Aldha1a^+^Vim^+^ EdCs. **(D-D”,F-F”)** Immunostaining for MHC (magenta) and α-SMA (white) on sections of cryoinjured ventricles from Ctrl (n=5 for *egr3* OE, n=6 for *Tg(hsp70l:Cre);egr3^flox/flox^*, D), *egr3* OE (D’), and *Tg(hsp70l:Cre);egr3^flox/flox^* (D”) zebrafish, and from *flt1^+/+^* (n=5, F) and *flt1^-/-^* (n=5, F’) zebrafish at 7 dpci. **(E,G)** Quantification of α-SMA^+^ cell number within the wound of the indicated genotypes at 7 dpci. **(H,H’)** Immunostaining for Aldh1a2 (magenta) and α-SMA (yellow) with DAPI (cyan) counterstaining on sections of cryoinjured *flt1^+/+^* (n=4, H) and *flt1^-/-^* (n=4, H’) ventricles at 7 dpci. Arrowheads point to endocardial-associated α-SMA^+^ cells. **(I)** Quantification of endocardial-associated α-SMA^+^ cell number within the wound on ventricle sections of the indicated genotypes at 7 dpci. White and orange dotted lines delineate the injured areas. White dotted boxes correspond to the magnified regions. Scale bars: 100 µm. Data show mean ± SEM (two-tailed, unpaired Student’s *t*-test with *p* values shown in the graphs).

EndoMT is a process by which endothelial cells lose their identity and give rise to mesenchymal cells such as fibroblasts and myofibroblasts, contributing to organ fibrosis, including cardiac fibrosis (Kovacic et al., 2019; Piera-Velazquez and Jimenez, 2019). Aberrant EndoMT leads to mammalian-like fibrosis after cardiac cryoinjury in zebrafish(Allanki et al., 2021). Myofibroblast differentiation is a hallmark of mammalian-like fibrosis post cardiac injury (Davis and Molkentin, 2014; Micallef et al., 2012). Notably, increased *Egr3* expression promotes pro-fibrotic responses in scleroderma and leads to myofibroblast accumulation in lesional dermis (Fang et al., 2013). To investigate whether Egr3 plays a role in myofibroblast differentiation during cardiac regeneration, we leveraged recently developed gain- and loss-of-function genetic tools (da Silva et al., 2024). Specifically, we utilized an *egr3* overexpression line *Tg(hsp70l:Gal4);Tg(UAS:egr3)*, abbreviated as *egr3*-OE, and an *egr3* floxed line crossed to an *hsp70l:Cre* line (Fig. S7A). We subjected these zebrafish along with their control siblings to cardiac cryoinjury and heat shock treatment and analyzed recombination efficiency and expression (Fig. S7B-D). To assess myofibroblast differentiation, we stained *egr3*-OE ventricles and controls for α-SMA expression at 7 dpci and found an increased number of α-SMA^+^ cells in the injured area following *egr3* OE (Fig. 4D,D’,E). Conversely, *Tg(hsp70l:Cre);egr3^flox/flox^*(abbreviated as *egr3^flox/flox^ + Cre*) and *Tg(hsp70l:Cre);egr3^flox/+^*(abbreviated as *egr3^flox/+^ + Cre*) ventricles displayed a significant decrease of α-SMA^+^ cells in the injured area compared with controls (Fig. 4D,D”,E, Fig. S7E-F), together suggesting a role for *egr3* in promoting myofibroblast differentiation.

Since *egr3* is downregulated in cryoinjured *flt1*^-/-^ hearts, we quantified the number of α-SMA positive cells in these ventricles and found a significant reduction (Fig. 4F-G), consistent with the reduced scarring observed in these animals. As *egr3* is upregulated in the regenerating endocardium, we quantified endocardial-associated α-SMA^+^ myofibroblasts, which we define as myofibroblasts co-localized with, in morphological continuity with, or in close proximity to EdCs. In agreement with the decrease in total α-SMA^+^ myofibroblasts within the injured tissue, endocardial-associated α-SMA^+^ myofibroblasts decreased by more than two-fold in cryoinjured *flt1^-/-^* hearts (Fig. 4H-I). Altogether, these data indicate that *flt1* mutant hearts exhibit a hyper-invasive endocardial phenotype as well as reduced endocardial-associated myofibroblast differentiation, the latter likely due to the downregulation of *egr3*.

Myofibroblasts are responsible for the production and deposition of extracellular matrix (ECM) proteins. ECM molecules can regulate cardiomyocyte mobilization and proliferation by directly signaling to cardiomyocytes or altering the microenvironment (Bassat et al., 2017; Chablais and Jazwinska, 2012; Chen et al., 2016; Fang et al., 2013; Koth et al., 2020; Notari et al., 2018; Wang et al., 2013; Wu et al., 2020). Previous studies have shown that ECM deposition strongly influences cardiomyocyte repopulation in injured zebrafish hearts (Allanki et al., 2021; Constanty et al., 2024; Wang et al., 2013). Therefore, we reasoned that alterations in *egr3* expression might affect cardiomyocyte repopulation following cardiac cryoinjury. To investigate this possibility, we quantified cardiomyocyte protrusions in cryoinjured ventricles at 7dpci and observed that they were significantly shorter in *egr3*-OE ventricles and significantly longer in *Tg(hsp70l:Cre);egr3^flox/flox^*(Fig. 5A-B) and *Tg(hsp70l:Cre);egr3^flox/+^* (Fig. S7G-H) ventricles compared with their respective control siblings. The number of cardiomyocyte protrusions remained similar across all groups (Fig. 5C, Fig. S7I). Cardiomyocyte protrusion length was also increased in *flt1* mutant ventricles at 7 dpci (Fig. 5D-E) while their number was not affected (Fig. 5F). Additionally, we noted a significant increase in cardiomyocyte proliferation in *Tg(hsp70l:Cre);egr3^flox/flox^*(Fig. 5G-H) and *Tg(hsp70l:Cre);egr3^flox/+^* ventricles (Fig. S7J-K), one similar to that observed in *flt1-/-* ventricles (Fig. 2C-D). Although cardiomyocyte proliferation in *egr3*-OE ventricles appeared reduced compared with control siblings, the difference was not statistically significant (Fig. S7L-M). To assess whether *egr3*-OE impacted scarring, we performed AFOG staining at 90 dpci. As expected from the previous results, scar area was significantly bigger in *egr3*-OE ventricles than in controls (Fig. 5I-K). These findings suggest that *egr3* exerts an inhibitory effect on cardiomyocyte repopulation, potentially by modulating myofibroblast-mediated ECM deposition, thereby altering the microenvironment for cardiomyocyte mobilization and proliferation. Collectively, our results indicate that *flt1* deletion restricts myofibroblast differentiation and promotes cardiomyocyte repopulation by downregulating endocardial *egr3*.

**Fig. 5.**
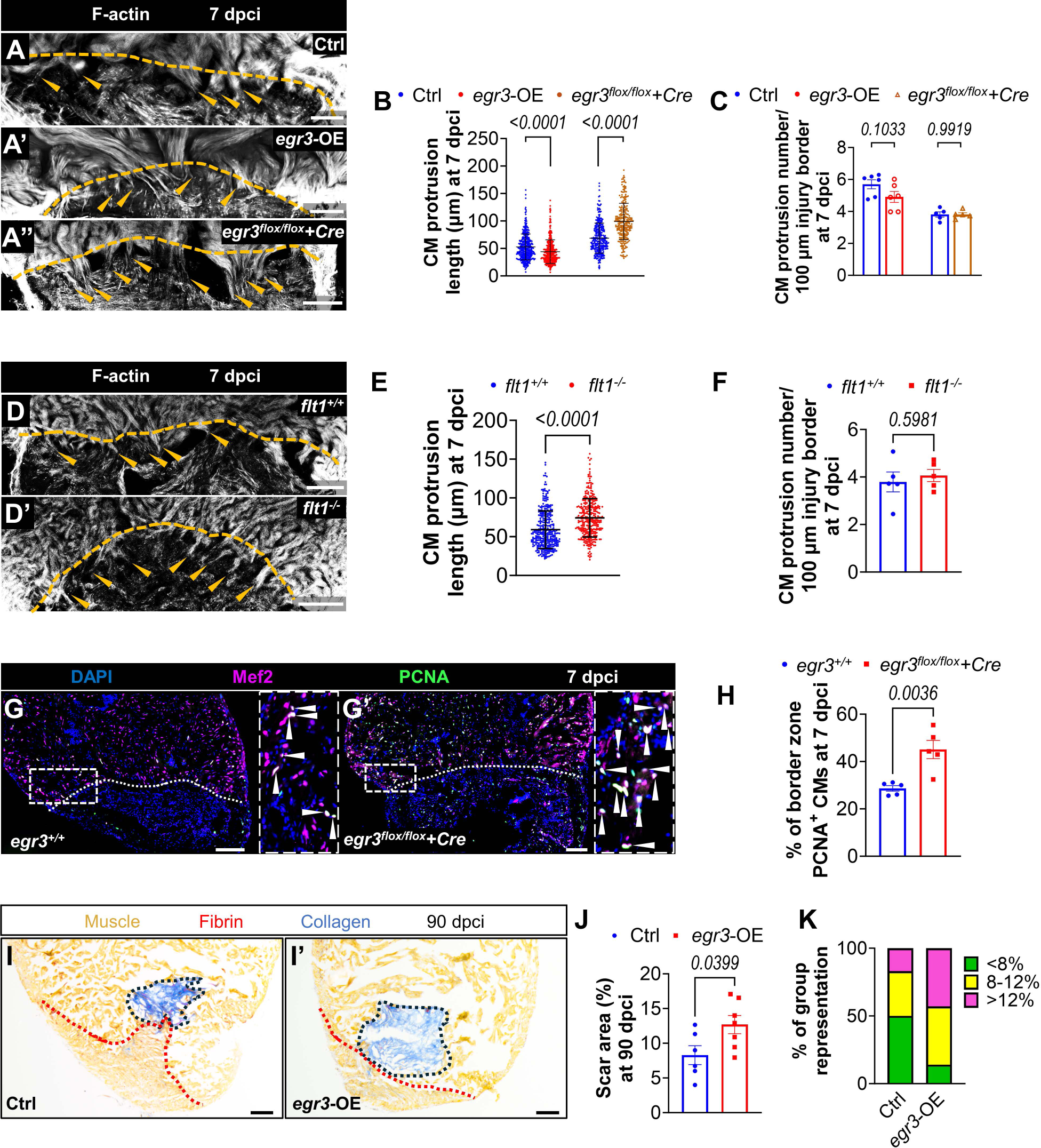
*flt1* deletion promotes cardiomyocyte repopulation by downregulating *egr3*. **(A-A”)** Phalloidin staining for F-actin (white) on 50-μm thick sections of cryoinjured ventricles from non-transgenic Ctrl (n=6 for *egr3* OE, n=5 for *Tg(hsp70l:Cre);egr3^flox/flox^*, A), *egr3* OE (n=6, A’), and *Tg(hsp70l:Cre);egr3^flox/flox^*(n=5, A”) zebrafish at 7 dpci. Orange arrowheads point to the protruding CMs in the injured area. **(B,C)** Quantification of CM protrusion length (B) and number (C) of the indicated genotypes at 7 dpci. **(D,D’)** Phalloidin staining for F-actin (white) on 50-μm thick sections of cryoinjured ventricles from *flt1^+/+^*(n=5, D) and *flt1^-/-^* (n=5, D’) zebrafish at 7 dpci. Orange arrowheads point to protruding CMs in the injured area. **(E,F)** Quantification of CM protrusion length (E) and number (F) in the indicated genotypes at 7 dpci. **(G,G’)** Immunostaining for Mef2 (magenta) and PCNA (green) with DAPI (blue) counterstaining on sections of cryoinjured ventricles from sibling *Tg(hsp70l:Cre);egr3^+/+^*(n=5, G) and *Tg(hsp70l:Cre);egr3^flox/flox^* (n=5, G’) zebrafish at 7 dpci. Arrowheads point to PCNA^+^ CMs. **(H)** Percentage of PCNA^+^ CMs in the border zone of the indicated genotypes at 7 dpci. **(I,I’)** AFOG staining on sections of cryoinjured non-transgenic Ctrl (n=6, I) and *egr3* OE (n=7, I’) ventricles at 90 dpci. Orange, muscle; red, fibrin; blue, collagen. Black and red dotted lines delineate scar and regenerated muscle wall areas, respectively. **(J)** Percentage of scar area relative to ventricular area in the indicated genotypes at 90 dpci. **(K)** Graphs showing the representation of groups of different scar area sizes in the indicated genotypes at 90 dpci. White and orange dotted lines delineate the injured areas. White dotted boxes correspond to the magnified regions. Scale bars: 100 µm. Data in (C,F,H,J) show mean ± SEM (two-tailed, unpaired Student’s *t*-test with *p* values shown in the graphs). Data in (B,E) show mean ± SD (two-tailed, Mann-Whitney *U* test with *p* values shown in the graphs).

## Discussion

Here we report that *flt1* inactivation enhances zebrafish cardiac regeneration by augmenting the endothelial response, consequently promoting cardiomyocyte regeneration, and limiting scarring.

FLT1 serves as a decoy receptor for VEGFA, VEGFB and placental growth factor (PlGF) (Ruiz de Almodovar et al., 2009). Previous studies in zebrafish embryos and larvae have shown that the *flt1* mutant endothelial phenotypes are Vegfaa dependent (Wild et al., 2017). Moreover, overexpression of *vegfba* and *plgf* has no impact on vessel sprouting (Klems et al., 2020). Here we used a mutated form of Vegfaa that has been shown to interfere with its affinity for Vegfr2 (Marin-Juez et al., 2016; Muller et al., 1997a; Muller et al., 1997b; Rossi et al., 2016) and found that the *flt1* mutant regeneration phenotypes were rescued, indicating increased Vegfa bioavailability in these animals.

Both coronary and endocardial regeneration have been shown to be critical to support tissue replenishment in regenerative and non-regenerative organisms (Apte et al., 2019; Das et al., 2019; DeBenedittis et al., 2022; Dube et al., 2017; Kivela et al., 2019; Lai et al., 2017; Miquerol et al., 2015; Tang et al., 2018). In addition, global deletion of *Vegfr1* at post-natal stages has been shown to increase angiogenesis in different vascular beds and reduce the infarct size in adult mice after ligation of the left anterior descending artery (Ho et al., 2012). Our results highlight the importance of revascularization for baseline regeneration and the requirement of an augmented endocardial response to enhance it. Moreover, the endocardium serves as a source of Vegfaa which regulates intraventricular coronary sprouting in a process termed coronary-endocardial anchoring (Karra et al., 2018; Marin-Juez et al., 2019). Coronary vessels form a vascular scaffold to guide cardiomyocyte development and regeneration (DeBenedittis et al., 2022; Marin-Juez et al., 2019). Therefore, enhancing coronary-endocardial anchoring might itself facilitate cardiomyocyte replenishment in *flt1* mutants.

Our data identified that MAPK/ERK signaling is increased in the regenerating endocardium in *flt1* mutants. MAPK/ERK signaling regulates cell proliferation, growth, and migration (Lavoie et al., 2020; Unal et al., 2017) and promotes Vegfa-dependent angiogenesis in zebrafish (Shin et al., 2016). *dusp6*, a phosphatase that inhibits ERK signaling (Marchetti et al., 2005; Muda et al., 1996), is expressed in cardiac endothelial cells after injury and *dusp6* zebrafish mutants display improved heart regeneration(Missinato et al., 2018). Moreover, stimulation of MAPK signaling has been shown to stimulate endothelial cell proliferation while inhibiting TGFβ induced EndoMT (Ichise et al., 2014). Thus, it is reasonable to hypothesize that increased Vegfa signaling in cryoinjured *flt1* mutant hearts limits endocardial EndoMT via increased MAPK/ERK activation. Indeed, we find reduced number of endocardial α-SMA^+^ myofibroblasts in injured *flt1* mutant ventricles, further supporting this possibility.

Myofibroblasts contribute to fibrosis by depositing ECM. Changes in ECM composition can influence cardiomyocyte regeneration. ECM-bound proteins can signal directly to cardiomyocytes and determine the physical properties of the microenvironment, impacting cardiomyocyte mobilization and division (Bassat et al., 2017; Chablais and Jazwinska, 2012; Chen et al., 2016; Fang et al., 2013; Koth et al., 2020; Notari et al., 2018; Wang et al., 2013; Wu et al., 2020). Fibrosis is the main cause of heart failure after MI in non-regenerative models. However, fibrosis also develops transiently in the regenerating zebrafish heart, where ablation of collagen producing cells impairs cardiomyocyte proliferation (Sanchez-Iranzo et al., 2018). By manipulating its expression, we show that Egr3 negatively regulates myocardial protrusion and proliferation, likely due to the effect of these manipulations on myofibroblast differentiation. It is interesting to note that *egr3* is upregulated as part of the natural regeneration program deployed by the cryoinjured zebrafish heart. However, its downregulation in conditions of increased Vegfa bioavailability leads to improved regeneration. *EGR3*/EGR3 expression is induced by TGFß signaling in normal skin fibroblasts (Fang et al., 2013). TGFß signaling is induced after cardiac injury in zebrafish and its inhibition is detrimental for heart regeneration, suggesting the need for transient fibrosis as part of a successful cardiac regeneration program (Chablais and Jazwinska, 2012, Sanchez-Iranzo et al., 2018). It is possible that by abrogating Egr3 expression, the effect of TGFß signaling activation is partially attenuated, limiting excessive myofibroblast differentiation and ECM deposition while conserving key components of the fibrotic response. These conditions might create a permissive microenvironment favoring regenerative processes such us cardiomyocyte dedifferentiation, protrusive behavior, and proliferation.

Overall, our data indicate that Vegfa-induced attenuation of the fibrotic response enhances cardiac regeneration, at least in part, by providing a more permissive milieu for cardiomyocyte replenishment.

## Acknowledgement

We thank R. Ramadass and K. Mattonet for help with microscopy, and T. Molina-Villa, P. Goumenaki, T.-L. Tseng, and Z.-F. Fang for discussions on the study and critical comments on the manuscript. We thank our animal house staff for their excellent support. We also thank M.T.M. Mommersteeg (University of Oxford) for sharing the pre-analyzed data of GSE138181 for reanalysis. Z.-Y.W. and Q.-C.W. were recipients of fellowships from Kerckhoff-Stiftung, China Scholarship Council, and Central South University. R. Marín-Juez is currently supported by an FRQS Junior-1 award.

## Author Contributions

Conceptualization: Z.-Y.W., D.Y.R.S and R.M.-J. Methodology: Z.-Y.W., D.Y.R.S and R.M.-J. Investigation: Z.-Y.W., A.M., Q.-C.W., A.R.-S., T.J., S.G., M.L., and J.D. Formal analysis: all authors. Writing - Original Draft: Z.-Y.W., D.Y.R.S and R.M.-J. Writing - Review and Editing: All authors. Supervision: D.Y.R.S and R.M.-J. Project administration: D.Y.R.S and R.M.-J. Funding acquisition: D.Y.R.S and R.M.-J.

## Data availability

RNA-seq data is deposited in NCBI GEO database under accession number GSE264406.

